# Salience network dynamics underlying successful resistance of temptation

**DOI:** 10.1101/129676

**Authors:** Rosa Steimke, Jason S. Nomi, Vince D Calhoun, Christine Stelzel, Lena M. Paschke, Robert Gaschler, Henrik Walter, Lucina Q. Uddin

## Abstract

Self-control and the ability to resist temptation are critical for successful completion of long-term goals. Contemporary models in cognitive neuroscience emphasize the primary role of prefrontal cognitive control networks in aligning behavior with such goals. Here we use gaze pattern analysis and dynamic functional connectivity fMRI data to explore how individual differences in the ability to resist temptation are related to intrinsic brain dynamics of the cognitive control and salience networks. Behaviorally, individuals exhibit greater gaze distance from target location (e.g. higher distractibility) during presentation of tempting erotic images compared with neutral images. Individuals whose intrinsic dynamic functional connectivity patterns gravitate towards configurations in which salience detection systems are less strongly coupled with visual systems resist tempting distractors more effectively. The ability to resist tempting distractors was not significantly related to intrinsic dynamics of the cognitive control network. These results suggest that susceptibility to temptation is governed in part by individual differences in salience network dynamics, and provide novel evidence for involvement of brain systems outside canonical cognitive control networks in contributing to individual differences in self-control.

## Introduction

In our daily lives we constantly encounter situations that evoke conflicting response tendencies: on the one hand impulsive reactions towards tempting stimuli and on the other hand actions that serve the realization of previously set goals (Hofmann et al., 2012). Self-control is correlated with well being (Hofmann et al., 2014), and self-control failure has been related to addiction, obesity, posttraumatic stress disorder, depression, and attention-deficit hyperactivity disorder (Bechara, 2005; Konttinen et al., 2009; Özdemir et al., 2014; Schweitzer & Sulzerazaroff, 1995; Walter et al., 2010). This is why it is of great scientific interest to understand why some individuals are able to resist when faced with temptation, while others fail.

Erotic and sensual images are powerful visual temptations. The advertisement industry frequently makes use of erotic images (Reichert & Carpenter, 2004) because they are very salient, and trigger us to involuntarily look towards them (Sennwald et al., 2016). This might be the case because they trigger evolutionarily meaningful attention allocation and approach behaviors (Fromberger et al., 2012). Here we investigate the neural basis for individual differences in self-control in the face of temptation using a combination of eyetracking and dynamic functional connectivity fMRI.

The most prominent model of self-control is the dual-systems approach, which assumes that a reflective system serving higher-level goal representations can exert control over an impulsive system that reacts to stimuli in a direct automatic manner (e.g., Hofmann et al., 2009; Metcalfe & Mischel, 1999; Strack & Deutsch, 2004). The reflective system has been mainly associated with fronto-parietal cognitive control networks, while the impulsive system has been linked with visceral and sensory regions (McClure & Bickel, 2014). Prefrontal cortical regions have been associated with self-control (e.g. Hare et al., 2009; Hayashi et al., 2013). Based on this model, the cognitive control network is a prime candidate for studying individual differences in self-control.

Another potential candidate for explaining individual differences in self-control is the salience network. The salience network is comprised of bilateral insula, dorsal anterior cingulate cortex (dACC) and other subcortical and limbic structures (Seeley et al., 2007), and is implicated in the direction of attention towards important stimuli and integration of top down appraisal and bottom up visceral and sensory information (see Uddin, 2015, for review). This central role in integrating information is reflected in its unique functional and structural connectivity profile. For example, the different insular nodes within the salience network are associated with distinct functional connectivity profiles; the dorsal anterior insular cortex coactivates with areas associated with cognitive processing, the ventral anterior insular coactivates with areas associated with affective processing and the posterior insular coactivates with sensorimotor processing areas (Chang et al., 2013; Uddin et al., 2014).

In the task presented here, attention allocation towards task relevant information is in conflict with attention allocation towards task irrelevant, yet intrinsically relevant, information (erotic distractors). As the salience network has been implicated in the allocation of attention processes towards task relevant information by interacting with other networks and the coordination of resources (Uddin, 2015), individual differences in salience network functioning might play an important role in explaining why some participants stay on task while others yield to the erotic distraction.

Most self-control research has focused on the down-regulation of impulses by cognitive control networks when examining individual differences in self-control, while studies of bottom-up processes that influence self-control are underrepresented (but see Steimke et al., 2016; Ludwig et al, 2013). Task based fMRI studies have indicated that self-control involves dorsolateral prefrontal cortex modulation of a value signal in the ventromedial prefrontal cortex (Hare et al., 2009). Recently, spontaneous fluctuations in resting state brain activity have been shown to demonstrate reproducible correlations across brain regions organized into networks (Shehzad et al., 2009). Because resting state networks are thought to represent individual differences in the brain’s functional organization, resting state fMRI has become a leading approach for understanding individual differences in behavior (Dubois & Adolphs, 2016).

Dynamic functional connectivity of resting state fMRI data is a new approach that accounts for the non-stationarity of brain signals and enables the study of brain dynamics underlying behavior. Whereas the static functional connectivity approach assumes that the connectivity pattern of the brain remains stable over time, the dynamic functional connectivity approach accounts for moment-to-moment variability in connectivity profiles. Within this framework the brain engages in reoccurring time-varying functional relations that can be referred to as functional connectivity “states” (Calhoun et al., 2014; Hutchison et al., 2013). By taking time variation into account, the dynamic approach can give a more nuanced understanding of brain connectivity, which is vital for understanding the source of individual differences. Our previous work examining the dynamic functional connectivity profile of different insular subregions found partially distinct and partially overlapping dynamic state profiles of the anterior, ventral and posterior insular subdivisions, highlighting aspects of salience network dynamics that have been previously overlooked (Nomi et al., 2016). Whole brain dynamic state characteristics are related to individual differences in executive function (Jia et al., 2014; Nomi et al., 2017; Yang et al., 2014) as well as mental illnesses including schizophrenia and bipolar affective disorder (Damaraju et al., 2014; Rashid et al., 2014). No previous studies have considered the relationship between salience network dynamics and self-control.

Here we present results of a study examining the relationship between brain network dynamics and self-control in the face of temptation. Broadly, we expected that susceptibility to interference from visual distraction would be reflected in brain network dynamics. We predicted that individual differences in self-control would be related to cognitive control network dynamics, salience network dynamics, or both. We explored these potential mechanisms underlying individual differences in self-control in a relatively large sample of 94 adults.

### Methods

#### Participants

Ninety-four current or former university or college students (Mean age = 25.93, SD = 3.84; 54 females) were included in the analysis. These data were part of a larger data set of 126 participants who also participated in additional fMRI tasks, self-control and cognitive control paradigms (Paschke et al., 2016; Sekutowicz et al., 2016). Of all participants, 109 had valid behavioral and eyetracking data (Steimke et al., 2016). Fifteen participants were excluded because either the fMRI registration process was not successful, or because the fMRI scans did not cover the whole brain.

#### Self-Control Task

In a task designed to assess self-control, participants were instructed to attend to a cued target location (left or right side of the screen) while facing the challenge to sustain attention despite neutral and erotic pictures presented as distractors on the other side of the screen (Figure 1 and 2; see Steimke et al., 2016 for behavioral and eyetracking results of the task). Eyetracking data were acquired using a video-based eyetracker (sampling rate: 250 Hz. spatial resolution: 0.05°, Cambridge Research Systems, UK). Participants were seated 36 cm from the screen. In order to reduce movement, participants were instructed to rest their chin and forehead on a chin rest. The distracting images were presented for a variable duration and elicited participants’ eye gaze to shift away from the cued target location, resulting in poorer performance on the task, which was to identify by button press whether a white target letter was an “E” or an “F”. Variable distractor durations were introduced to prevent participants from anticipating when the target letter would be presented. Motivation of participants to perform accurately on the task was enhanced by offering the chance of a 10 Euro reward for accurate performance on one trial, which was randomly selected after completion of the task. Target letters were presented for 10 ms against a dark grey background, and participants were instructed to respond as quickly and accurately as possible. The distractors consisted of neutral pictures (eg. neutrally rated objects or scenes) and erotic pictures (pictures of couples in erotic situations) displayed on the contralateral side of the screen relative to the target location. Note that the task also included other conditions not reported here. Pictures were selected on the basis of valence, arousal, and attraction ratings from 96 independent participants and erotic and neutral pictures were matched for brightness and complexity. Eye gaze distance from target location was used as a dependent variable. Specifically, the gaze distance difference score between trials with erotic and trials with neutral distraction was used. Gaze distance was used, as it is the most direct measure distractibility. Additionally, it was found that gaze distance, not reaction time, was related to delay of gratification in this task: participants who couldn’t resist to eat one sweet immediately instead of waiting for two sweets after 45 minutes also showed greater gaze distance from target when erotic pictures were presented than participants who chose to wait (Steimke et al., 2016). A short test of fluid intelligence (LPS, Leistungsprüfsystem Unterteil 3, Horn, 1983) was performed in order to be able to control for individual differences in intelligence.

**Figure 1.**
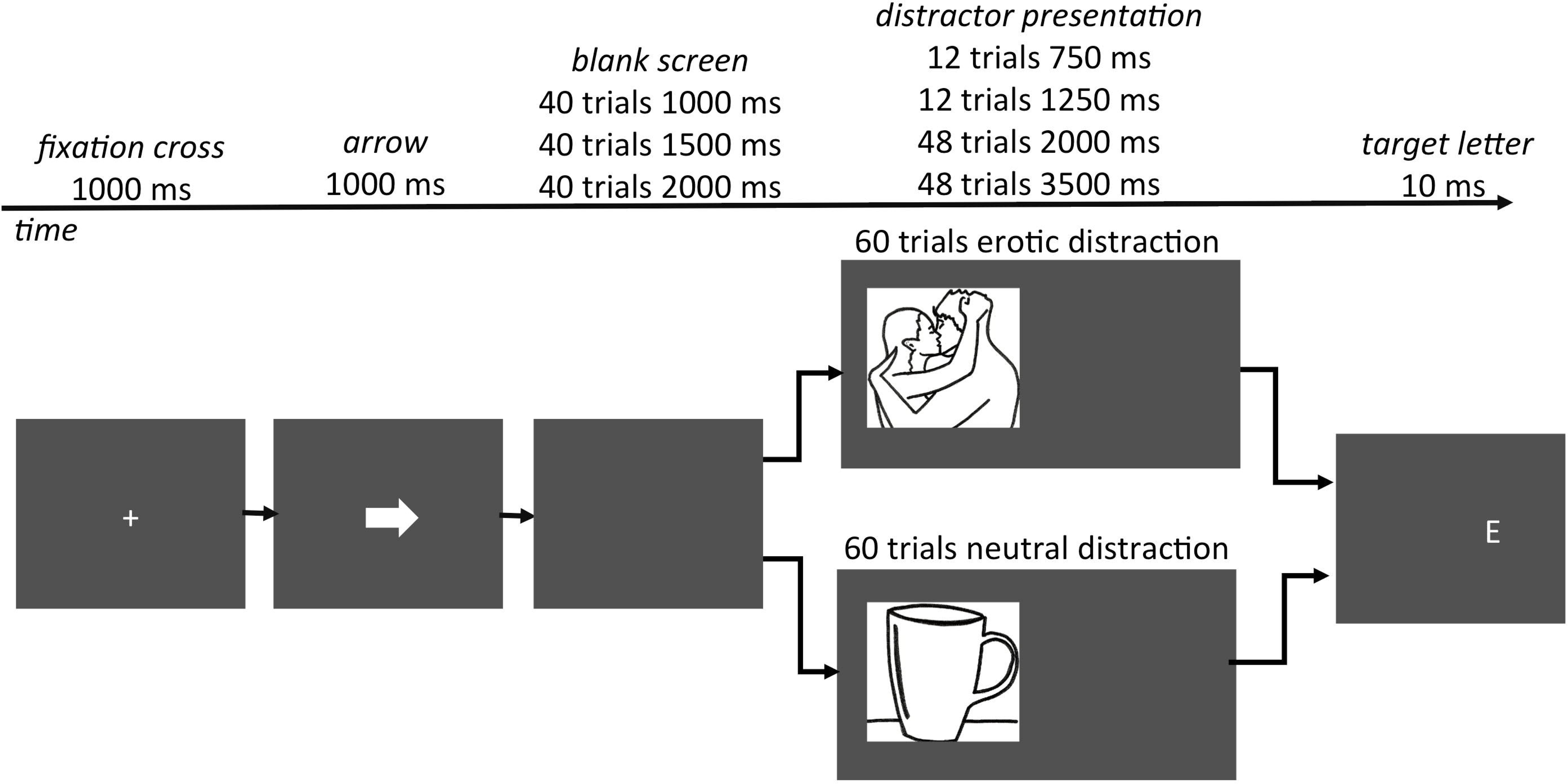
Display of the trial timing. A trial starts with a fixation cross. The fixation cross is followed by an arrow indicating the location of the next target letter “E” or “F” presented 5.9° of visual angle left or right from the center. After the arrow presentation the screen a cleared screen is presented for variable delay. Afterwards either a neutral or an erotic distractor is presented for a variable duration immediately followed by the target letter. Drawings are placeholders for photographs from the International Affective Picture System (Lang et al., 2008) and the internet.

**Figure 2.**
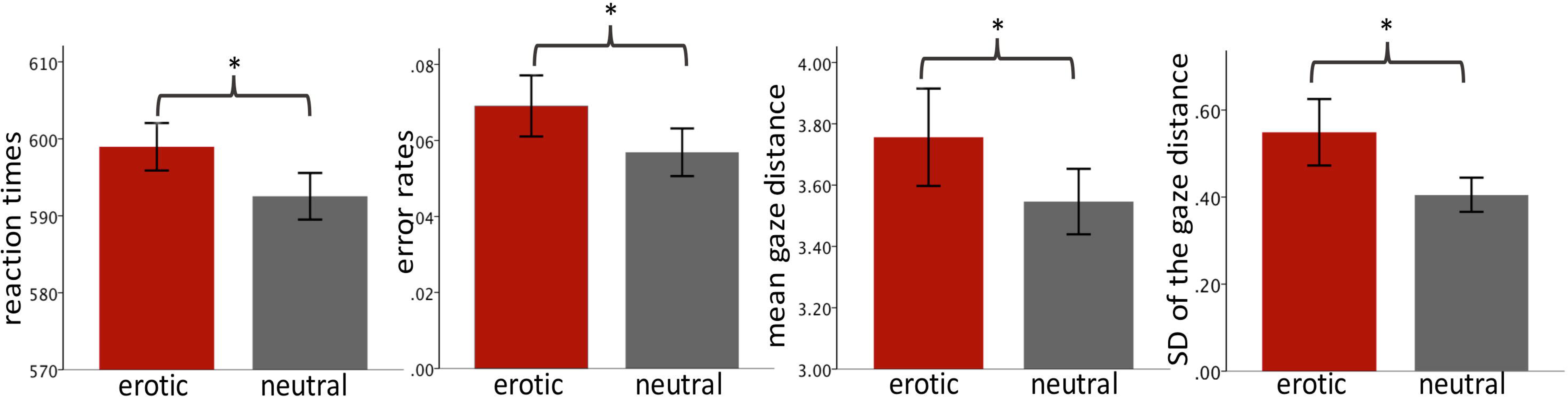
Display of behavioral and eyetracking data. Mean reaction times, percent errors, gaze distance and standard deviation of the gaze distance (SD). Asterisks (*) indicate a significant difference at *p* < 0.05. Error bars represent the 95% confidence interval for within-subjects comparisons (Loftus and Masson, 1994).

#### FMRI data acquisition and preprocessing

Each participant underwent resting-state fMRI scans during which they kept their eyes open and fixated on a fixation cross. Whole-brain fMRI data were collected with a 3 Tesla Siemens Tim Trio MRI scanner (Siemens, Erlangen, Germany) on a separate day from the behavioral testing. Using a 12 channel head coil, 32 slices were acquired in descending order with a T2*-sensitive one-shot gradient echo echo-planar sequence. To minimize motion, the head was fixated with cushions. The following parameters were used: repetition time of 2 s, echo time of 25 ms, flip angle of 78, data acquisition matrix of 64x64, field of view of 24 cm, voxel size of 3x3x3mm and an inter-slice gap of 0.75 mm, 180 volumes. Preprocessing was performed with the Data Processing Assistant for Resting-state fMRI (http://restfmri.net). In order to ensure data were at signal equilibrium, the first 10 volumes were removed. Slice time correction was performed, the data were realigned, normalized to an EPI template, and smoothed with a 8mm Gaussian kernel.

### Independent Component Analysis

The first step in the dynamic functional network connectivity (dFNC) analysis was to parcellate the brain into regions of interest using a high-model order group independent component analysis (ICA) implemented with the GIFT toolbox (http://mialab.mrn.org/software/gift/) using the infomax algorithm (Calhoun, Adali, Pearlson, & Pekar, 2001; Calhoun & Adali, 2012). A high-model order of 100 independent components was chosen based on previous work demonstrating that this number of components sufficiently parcellates major brain networks (default mode network: DMN; cognitive control network: CCN; salience network: SN) into individual brain areas that allows for more fine grained examination of network node interactions (Damaraju et al., 2014; Nomi et al., 2016, 2017). Additional research demonstrates that model orders of 100 and below have better reproducibility than model orders higher than 100 (Abou-Elseoud et al., 2010). Stability of ICs were ensured by repeating the infomax algorithm 10 times using ICASSO and selecting the central run for further analysis. Subject specific spatial maps and time courses were back-reconstructed using the GICA1 method (Erhardt et al., 2011).

The ICA produced 100 ICs that were then subjected to visual inspection in order to eliminate components containing white matter, cerebral spinal fluid, movement, or large amounts of high-frequency information (Allen et al., 2011; Damoiseaux et al., 2006; Nomi et al., 2016, 2017). This resulted in 44 ICs of interest that were grouped into functional networks by visual inspection following previous work: SN, CCN, visual network, components of the temporal lobes, DMN, components of the cerebellum, and a subcortical network (see Figure 3a). The correlation matrix representing overall strength of coupling between these ICs can be seen in Figure 3b.

**Figure 3.**
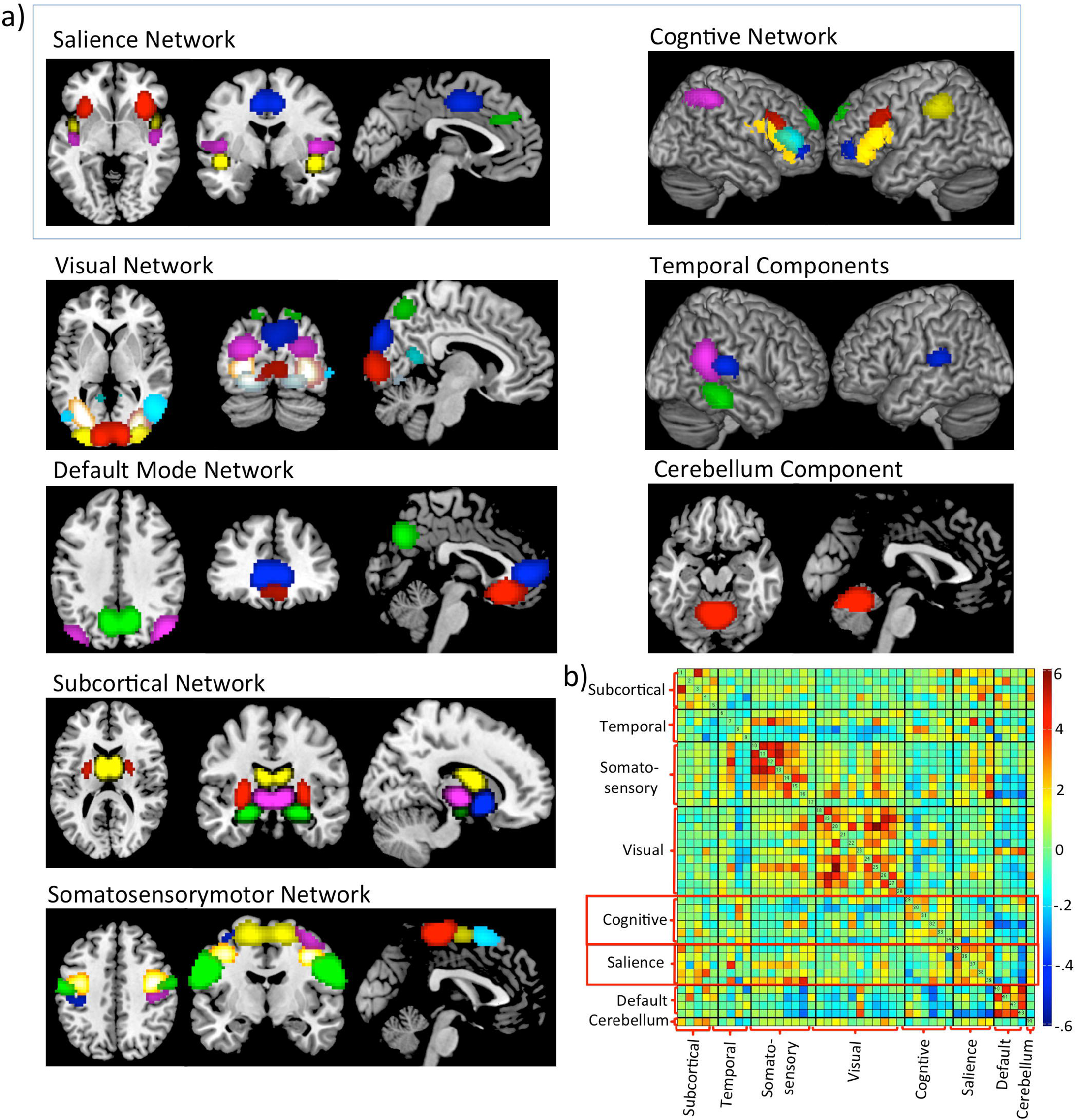
Brain networks and Static Connectivity. a) Display of the nodes identified by the ICA grouped into functional networks; each color represents a node within the network. b) Static whole brain functional connectivity correlation matrix; cognitive and salience networks used in the dynamic resting state analysis are highlighted by red boxes.

#### Independent Component Post-processing

Post-processing of non-noise ICs in GIFT consisted of despiking, detrending (linear, cubic, quadratic), regression of the Friston 24 motion parameters, and a low pass filter (0.15 Hz). Despiking replaces outliers in IC time courses larger than the absolute median deviation with a third-order spline fit to clean portions of the data using AFNI’s 3dDespike algorithm. Despiking decreases the temporal derivative (DVARS) (Power et al., 2011) over IC time courses and eliminates artifacts in dFNC analyses (Damaraju et al., 2014).

#### Sliding Window Analysis

Post-processed IC time courses were analyzed by using a sliding window dFNC algorithm in GIFT using window sizes of 22 TRs (44 seconds) slid in 1 TR. A window size of 44 seconds was chosen as previous dFNC work utilized window sizes of 44 (Yang et al., 2014) and 45 seconds (Damaraju et al., 2014; Nomi et al., 2016). Furthermore, previous dFNC research has demonstrated that window sizes of 30-60 seconds capture distinct dynamic functional connections not found in larger window sizes (Damaraju et al., 2014; Hutchison et al., 2013), methodological dFNC work has shown that such window sizes represent real fluctuations in functional connectivity (Leonardi & Van De Ville, 2015; Sakoglu et al., 2010). Additional empirical research demonstrates that these window sizes are able to capture cognitive states (Shirer, Ryali, Rykhlevskaia, Menon, & Greicius, 2012; Wilson et al., 2015).

A tapered window consisting of a rectangle convolved with a Gaussian (σ = 3) was utilized in order to account for the limited number of time points in each sliding window. This produced a covariance matrix with the dimensions of 946 (sliding windows) x 148 (paired connections) per subject. In order to further account for noise that may arise from a limited number of time points, each covariance matrix was regularized using the graphical LASSO method (L1 norm) (Friedman, Hastie, & Tibshirani, 2008) of the inverse covariance matrix resulting in a correlation matrix (Damaraju et al., 2014).

#### Cognitive Control Network and Salience Network Dynamic States

Windowed correlation matrices for components within the cognitive control network and salience network were extracted and subjected to k-means clustering independently. The salience network consisted of the bilateral insular, the dorsal anterior cingulate cortex and the orbitofrontal cortex; the cognitive control network consisted of bilateral dorsolateral prefrontal and parietal regions (see Figure 3a for salience and cognitive control network nodes and Figure 3b for the extracted matrices). In order to be able to perform k-means clustering, the sliding windows of all participants were concatenated for the cognitive control and salience network separately. Using these concatenated salience and concatenated cognitive control networks, the number of optimal number of clusters was determined by using the elbow criterion applied to the cluster validity index derived from k-means clustering using “city block” distance function (Allen et al., 2014) performed for clustering values between 2-20. This analysis revealed that 5 is the optimal number of clusters for the salience network as well as for the cognitive control network, therefore the 5 cluster solutions for all results are presented.

For each of the 5 cognitive and the 5 salience network states, frequency and dwell time were calculated. Frequency was calculated as the percent that a brain state prevailed throughout the duration of the scan. Dwell time was calculated as the average length, measured in number of sliding windows, that a participant stayed in a given brain state. Pearson correlations were calculated to relate frequency and dwell time of cognitive and salience network states to distractibility by erotic temptation.

### Results

#### Dynamic Salience Network States

We identified 5 different salience network states (see Figure 4a). Note that the states are sorted by the average percent of time participants spent in each of the five states. On average, participants spent 39.53 percent of their time in state 1 (SD = 27.54), 16.55 percent of their time in state *2 (SD* = 16.59), 16.02 percent of their time in state 3 (SD = 15.72), 15.53 of their time in state 4 (SD = 17.06), and 12.37 percent of the time is state 5 (SD = 16.15). Repeating of k-means clustering to a total of 5 estimates revealed stability of the results (see Table 1). Considering the frequency spent in each state, there is a floor effect for some participants, meaning they did not spend any time in that state at all. Five participants did not enter into state 1, 21 participants did not enter into state 2, 22 participants did not enter into state 3, 19 did not enter into state 4, and 33 did not enter into state 5. Not all participants enter into each state because k-means clustering of the concatenated data matrix including all subjects allows for the possibility that individual subjects will not contribute to each state (Damaraju et al., 2015; Nomi et al., 2016, 2017) The dwell time on average was 22.56 (SD = 26.00) for state 1, 10.87 (SD = 8.97 for state 2, 9.50 (SD = 8.25) for state 3, 9.70 (SD = 9.53) for state 4, and 9.98 (SD = 12.45) for state 5.

**Figure 4.**
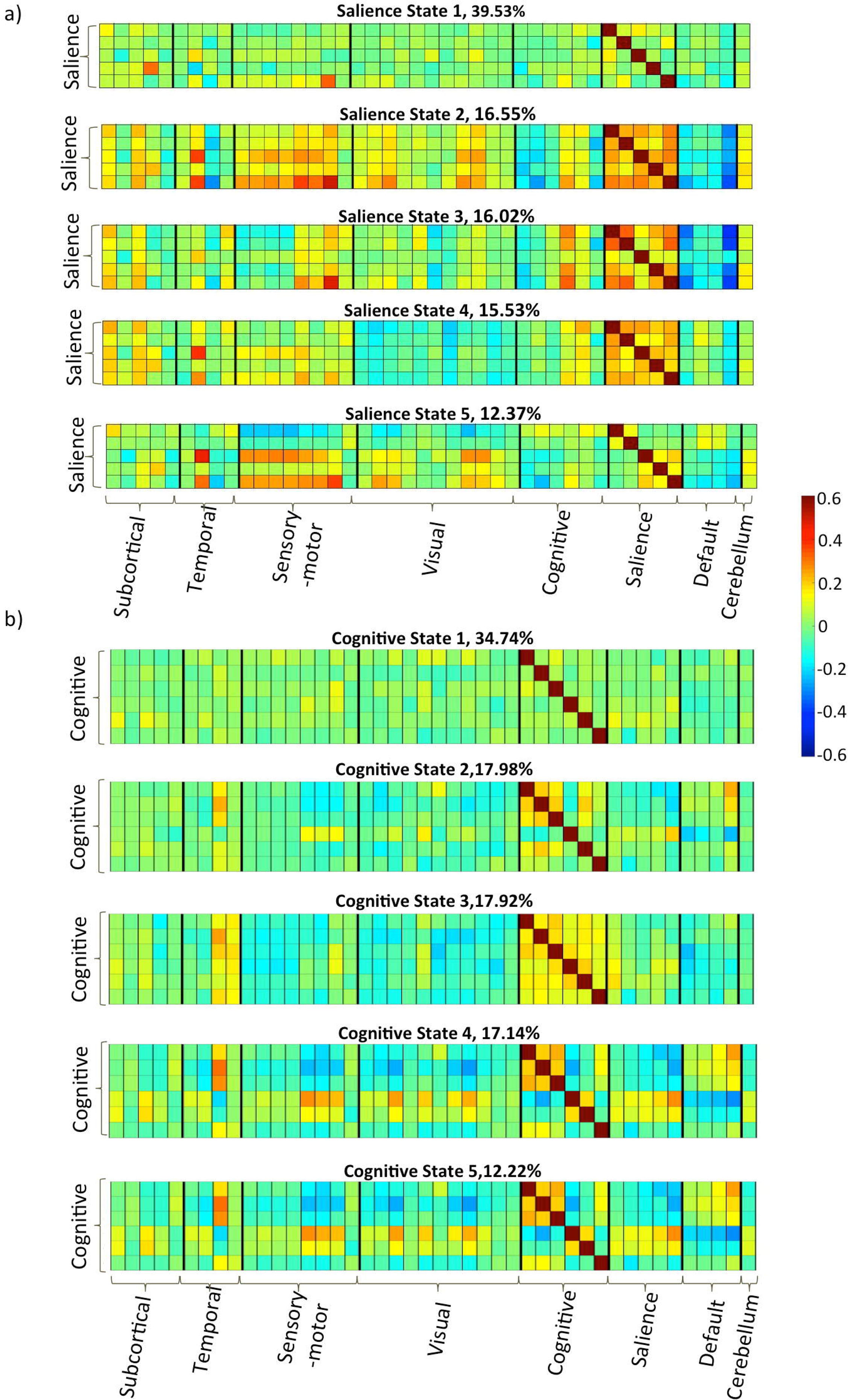
Display of the connectivity matrices of the five dynamic a) salience and b) cognitive control network states. States are order by the most frequent (state 1) to the least frequent (state 5). The frequency is indicated in percent time spent in each state.

**Table 1.** Percent of time spent in each salience network state. Repetition of k-means clustering reveals similar results, suggesting that the clustering is stable in this dataset.

|  | State 1 | State 2 | State 3 | State 4 | State 5 |
| --- | --- | --- | --- | --- | --- |
| **1<sup>st</sup> analysis** | 39.53% | 16.55% | 16.02% | 15.53% | 12.37% |
| **2<sup>nd</sup> analysis** | 39.50% | 16.50% | 16.04% | 15.52% | 12.44% |
| **3<sup>rd</sup> analysis** | 39.53% | 16.54% | 16.03% | 15.53% | 12.36% |
| **4<sup>th</sup> analysis** | 39.53% | 16.54% | 16.01% | 15.53% | 12.38% |
| **5<sup>th</sup> analysis** | 39.53% | 16.55% | 16.02% | 15.53% | 12.37% |

#### Dynamic Cognitive Control Network States

We identified 5 different cognitive control network states (see Figure 4b). The states are sorted by the average percent of time participants spent in each of the five states. On average, the participants spent 34.74 percent in state 1 (SD = 26.21), 17.98 percent of their time in state 2 (SD = 17.52), 17.92 percent in state 3 (SD = 17.67), 17.14 percent in state 4 (SD = 17.97), and 12.22 percent of their time in state 5 (SD = 13.84). Repeating of k-means clustering to a total of 5 estimates revealed stability of the results (see Table 2). Concerning the frequency, 3 participants did not adopt state 1, 20 participants did not adopt state 2, 18 participants did not adopt state 3, 18 did not adopt state 4, and 28 did not adopt state 5. The dwell time on average was 20.65 (SD = 26.37) for state 1, 10.96 (SD = 10.52) for state 2, 10.93 (SD =8.35) for state 3, 11.90 (SD =11.37) for state 4, and 10.76 (SD =11.19) for state 5.

**Table 2.** Percent of time spent in each cognitive control network state. Repetition of k-means clustering reveals similar results, suggesting that the clustering is stable in this dataset.

|  | State 1 | State 2 | State 3 | State 4 | State 5 |
| --- | --- | --- | --- | --- | --- |
| **1<sup>st</sup> analysis** | 34.74% | 17.98% | 17.92% | 17.14% | 12.22% |
| **2<sup>nd</sup> analysis** | 34.74% | 17.98% | 17.92% | 17.14% | 12.22% |
| **3<sup>rd</sup> analysis** | 34.55% | 17.97% | 18.21% | 17.13% | 12.15% |
| **4<sup>th</sup> analysis** | 34.61% | 17.75% | 18.07% | 17.10% | 12.46% |
| **5<sup>th</sup> analysis** | 34.74% | 17.98% | 17.92% | 17.14% | 12.22% |

#### Correlations with Eye Gaze Behavior

As reported in the behavioral paper (Steimke et al., 2016) distracting images elicited participants’ eye gaze to shift away from the target location, resulting in poorer performance on the task (Figure 2). Difficulty resisting temptations as indicated by gaze distance difference between erotic and neutral distractors was negatively correlated with the time spent in salience network state 4, *r(92)* = -0.26, *p* = 0.012. This correlation remained significant when excluding participants who did not spend any time in state 4, *r(73)* = -0.25, *p* = 0.031. Comparing participants who did not adopt state 4 at all with those who did ever adopt state 4, using between group t-testing, revealed marginally significant higher distraction by temptation for participants who adopted state 4, *t(91)* = 1.98, *p* = .051. State 4 represents a dynamic functional connectivity state wherein the salience network was negatively correlated with the visual network. All other states were not significantly correlated with performance on the temptation task (see Figure 5a). Posthoc analysis revealed that time spent in state 4 was negatively correlated with time spent in state 1 (*r(92)* = -0.36, *p* < 0.001). Further posthoc testing revealed that the significant negative correlation between time spent in state 4 and ability to resist temptation remained significant when controlling for age, gender and fluid intelligence as measured by LPS (*r* = -0.276, *p* = 0.008, *df* = 89). Correlating dwell time of the 5 salience network states with the ability to resist temptations revealed the same pattern as for frequency: state 4 showed a significant negative correlation, while the others did not (Figure 5b). There was no correlation between frequency or dwell time of any of the five cognitive control network states with self-control in the face of temptation (see Figure 6a and 6b).

**Figure 5.**
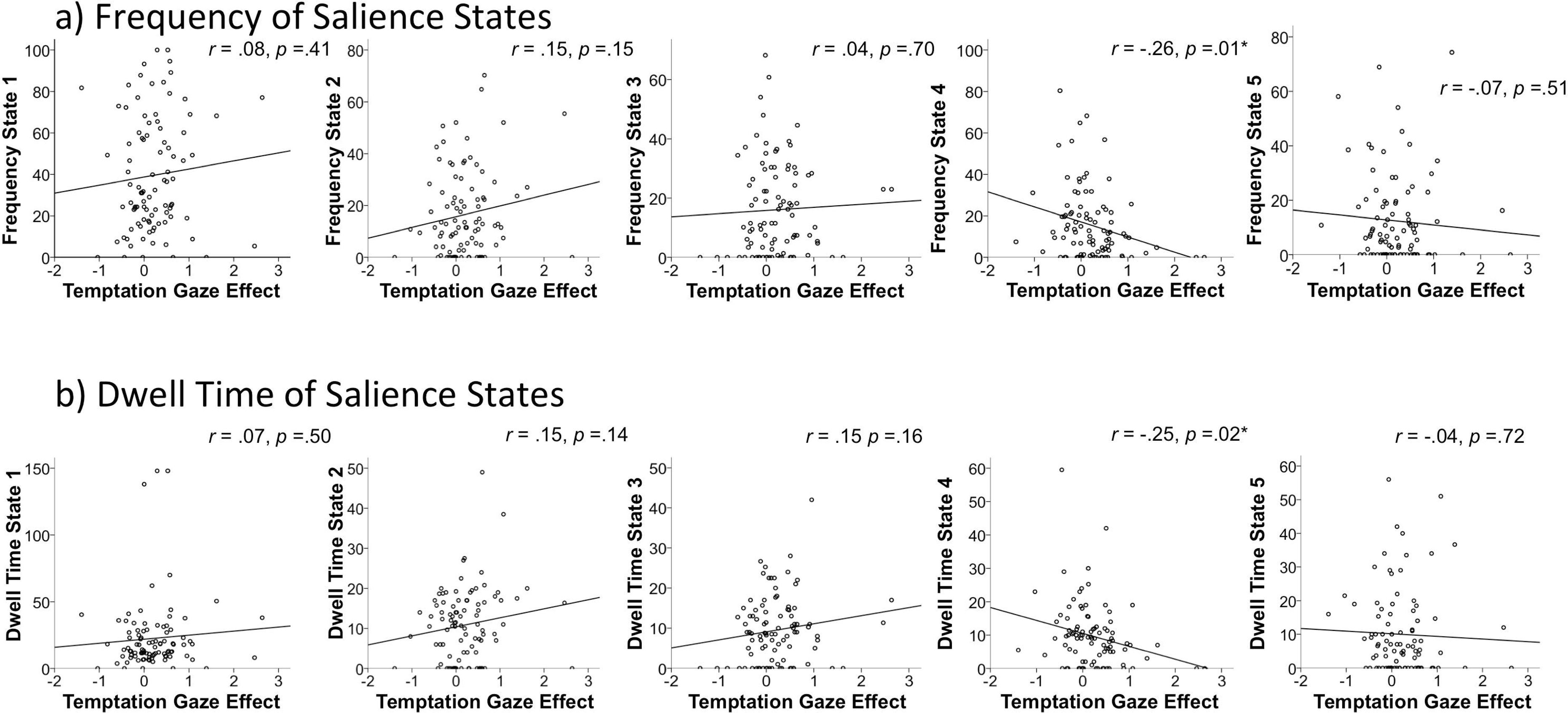
Correlations of a) frequency and b) dwell time of the 5 salience network states with the temptation gaze effect: the higher the temptation gaze effect, the more participants’ gaze drifted from the target location in the face of tempting distractors. The asterisk (*) indicates a significant difference at *p* < .05.

**Figure 6.**
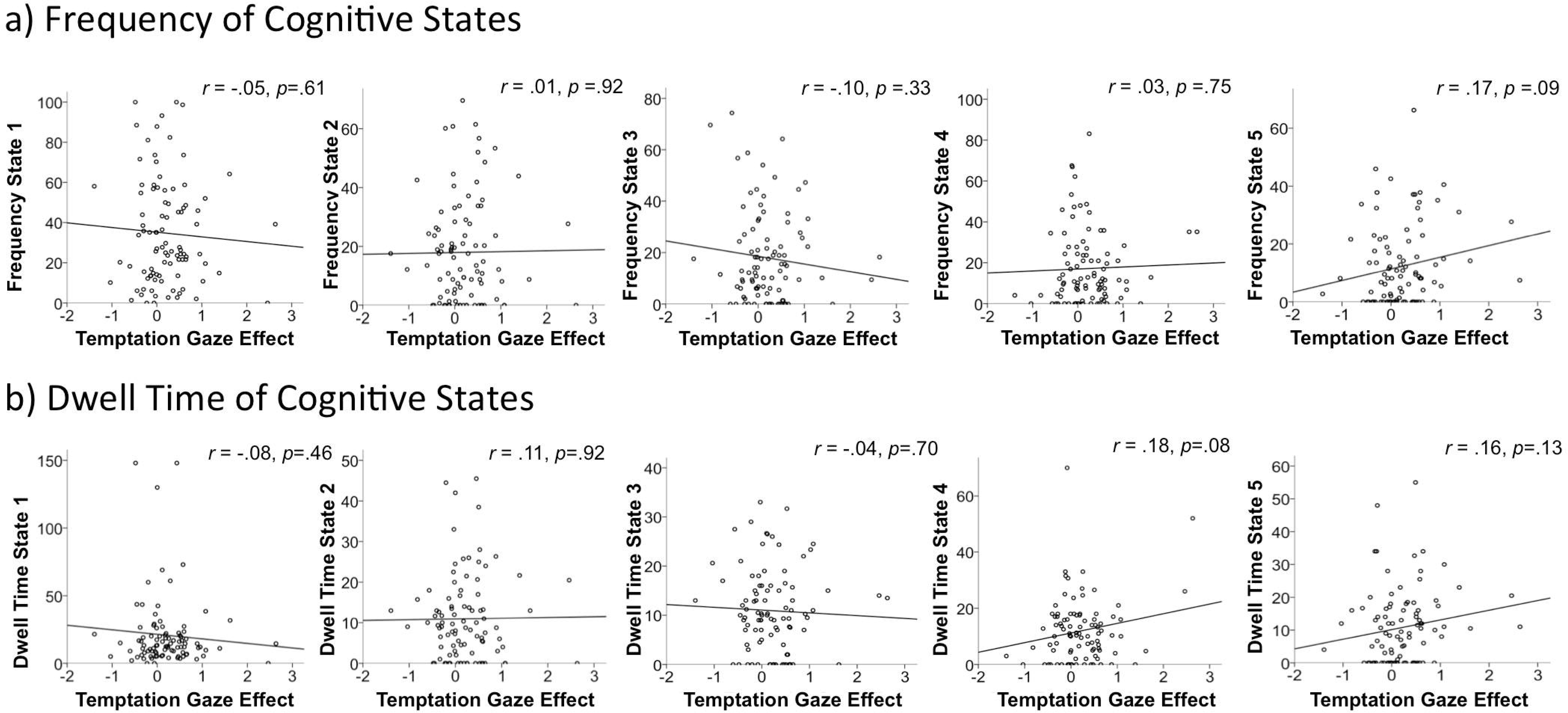
Correlations of a) frequency and b) dwell time of the 5 cognitive control network states with the temptation gaze effect: the higher the temptation gaze effect, the higher the distractibility by erotic images.

## Discussion

Self-control is critical for successful long-term goal attainment. Here we use gaze pattern analysis in a self-control task and dFNC analysis of resting state fMRI data to explore how individual differences in the ability to resist tempting distractors are related to intrinsic brain dynamics. We show that participants whose intrinsic connectivity patterns gravitate towards configurations in which salience detection systems are less strongly coupled with visual systems could resist tempting distractors more effectively.

Our results suggest that individuals whose brains spend more time in a state where salience and visual networks are decoupled were less distractible by erotic pictures. Most models of self-control posit a key role for prefrontal cognitive control networks in regulatory processes involved in overcoming the impulse to engage with salient distracting stimuli (Hare et al., 2009; Hayashi et al., 2013). The current results, in contrast, demonstrate for the first time that salience network dynamic coupling tendencies may contribute to individual differences in the ability to resist temptation.

The salience network, with key nodes in insular and anterior cingulate cortices, plays a central role in detection of behaviorally relevant stimuli and the coordination of neural resources. In particular, the dorsal anterior insular node of the salience network is thought to causally influence task-positive and default mode networks (Uddin et al., 2011). Salience network dysfunction has been linked with host of psychiatric conditions, particular those involving self-regulation and executive function deficits (Uddin, 2015). For these reasons, we predicted that individual differences in intrinsic salience network dynamics may contribute to the ability to focus and maintain attention when faced with tempting distractors.

Examination of salience network dynamics revealed the existence of 5 distinct connectivity states of this network (Figure 4). State 1, which was occupied nearly 40% of the time, was characterized by a large amount of correlations centered around zero. This is in line with previous studies (Damaraju et al., 2014; Nomi et al., 2017) which consistently show that brain states with the highest frequency of occurrence show a greater amount of correlations centered around zero compared to less frequently occurring states. State 2 was characterized by positive correlations of the salience network with the sensory motor network. Both State 2 and State 3 showed positive intercorrelations within the salience network and negative correlation of the salience with the default mode network. State 4 also showed a negative correlation of salience network with default network and intercorrelation within salience network. The most pronounced characteristic by which state 4 differed from the other states was the negative correlation of the salience network and the visual network. The least frequent state 5 showed mixed positive and negative correlation with regions in the sensorimotor networks, within salience network correlations centered around zero and correlation of salience network with default mode centered around zero.

Examination of cognitive control network dynamics revealed 5 cognitive control network states. The most frequent cognitive control network state (state 1) was occupied approximately 35 percent of the time and was characterized by a greater number of correlations centered around zero, in contrast to the other 4 states. This was the case for within cognitive control network intercorrelations and correlations of the cognitive control network with the rest of the brain.

The only significant relationship between brain network dynamics and individual differences in behavior was observed for salience network state 4. Both frequency and dwell time were significantly correlated with distractibility by erotic images as measured by gaze distance from target presentation.

In a recent whole brain dynamic functional connectivity study by Nomi et al. (2017), successful executive function was associated with spending more time in the most frequently occuring state, whereas in our study self-control was not associated with time spent in the most frequently occurring dynamic state. These results demonstrate for the first time how highly salient distractors can interfere with top-down control processes to a greater extent in individuals who exhibit specific patterns of intrinsic functional connectivity dynamics.

The current results provide more nuanced tests of the dual-systems model, which pits cognitive control systems against “impulsive” brain systems that react automatically to salient stimuli. Our separate analyses of cognitive control network dynamics and salience network dynamics indeed do not support the predictions of a traditional dual-systems approach. Instead, our findings suggest that when visual input has less access to salience detection systems, tempting erotic distractors are easier to ignore. These findings highlight the importance of considering “neural context” in studies of brain function; the idea that the functional relevance of a brain system depends on the status of other connected areas (McIntosh, 2004).

### Limitations

K-means is a powerful algorithm to identify a predefined number of clusters in a dataset.

However, it has also been criticized as the starting point of the algorithm can influence the clustering. If the number of clusters is chosen that does not match the dataset, the clustering can become unstable and repetition of the analysis can yield significantly different results. To address this limitation, we repeated the analysis 5 times. In Table 1 the percent time spent in each state for the five analyses is presented. The divergence between analysis lies below 0.1 percent, suggesting that the divergence between analyses is minor and the k-means clustering yielded stable results in our dataset.

With a p-value of 0.012, the results presented here do not survive the conservative Bonferroni correction for multiple comparisons. A corrected value would be marginally significant (p= .06). Therefore the results presented here should be regarded with caution and should be replicated in future studies. With a sample size of 94 we have a relatively large sample for a combined fMRI and eyetracking study. However, as was noted by Schönbrodt and Perugini (2013) a sample size of higher than 150-250 is even more reliable for examining correlations. Future work with larger sample sizes is needed to further support the results presented here.

### Conclusions

Studying salience network dynamics might deliver valuable insight into the origin of individual differences in self-control ability. We show that participants who spent more time in a brain network configuration in which salience detection systems are decoupled from visual systems could resist tempting distractors more effectively. This suggests that individual differences in self-control in the face of temptation might be driven in part by salience network functional connectivity context.

## Acknowledgements

We thank the Berlin School of Mind and Brain, the Humboldt University Berlin, and the Collaborative Research Centre “Volition and Cognitive Control” (DFG grant SFB 940/1 2013), Technical University Dresden, for financially supporting the project. This work was also supported by award R01MH107549 from the National Institute of Mental Health to LQU.

